# EZcalcium: Open Source Toolbox for Analysis of Calcium Imaging Data

**DOI:** 10.1101/2020.01.02.893198

**Authors:** Daniel A. Cantu, Bo Wang, Michael W. Gongwer, Cynthia X. He, Anubhuti Goel, Anand Suresh, Nazim Kourdougli, Erica D. Arroyo, William Zeiger, Carlos Portera-Cailliau

## Abstract

Fluorescence calcium imaging using a range of microscopy approaches, such as 2-photon excitation or head-mounted ‘miniscopes’, is one of the preferred methods to record neuronal activity and glial signals in various experimental settings, including acute brain slices, brain organoids, and behaving animals. Because changes in the fluorescence intensity of genetically encoded or chemical calcium indicators correlate with action potential firing in neurons, data analysis is based on inferring such spiking from changes in pixel intensity values across time within different regions of interest. However, the algorithms necessary to extract biologically relevant information from these fluorescent signals are complex and require significant expertise in programming to develop robust analysis pipelines. For decades, the only way to perform these analyses was for individual laboratories to write their own custom code. These routines were typically not well annotated and lacked intuitive graphical user interfaces (GUIs), which made it difficult for scientists in other laboratories to adopt them. Although the panorama is changing with recent tools like *CaImAn*, *Suite2P* and others, there is still a barrier for many laboratories to adopt these packages, especially for potential users without sophisticated programming skills. As 2-photon microscopes are becoming increasingly affordable, the bottleneck is no longer the hardware, but the software used to analyze the calcium data in an optimal manner and consistently across different groups. We addressed this unmet need by incorporating recent software solutions for motion correction, segmentation, signal extraction and deconvolution of calcium imaging data into an open-source, easy to use, GUI-based, intuitive and automated data analysis software, which we named *EZcalcium*.

## 1 Introduction

To understand brain function, neuroscientists are increasingly turning to synthetic and genetically encoded fluorescence indicators (calcium and voltage sensors) for monitoring dynamic fluorescence signals produced by neurons and glia (Lin and Schnitzer, 2016; Yang and Yuste, 2017). In particular, calcium imaging with 2-photon microscopy (Mostany et al., 2015; Svoboda and Yasuda, 2006) and micro-endoscopy with miniscopes or fiber photometry (Aharoni et al., 2019; Ghosh et al., 2011; Jacob et al., 2018; Liberti et al., 2017; Zhang et al., 2019) have provided key insights, such as how developing cortical networks undergo drastic transitions (Golshani et al., 2009; Rochefort et al., 2009) or how large neuronal ensembles encode spatial navigation (Dombeck et al., 2010). Calcium imaging offers several distinct advantages over traditional electrode recording techniques (Grewe and Helmchen, 2009; Grienberger and Konnerth, 2012): 1) it can be combined with genetic approaches (e.g., Cre-Lox) to probe neuronal activity in specific sub-populations of neurons (Goel et al., 2018; Yaeger et al., 2019) and glia (Srinivasan et al., 2016), either in specific subcellular compartments (e.g., axon boutons, dendritic spines, glial microdomains) (Broussard et al., 2018; Cichon and Gan, 2015) or in specific brain regions or cortical layers (Lacefield et al., 2019); 2) recordings can be carried out over periods of days or even weeks (Chen et al., 2013; He et al., 2018); 3) recordings can be made simultaneously in a large population of hundreds or thousands of neurons in multiple brain regions (e.g., an entire sub-network) (Sofroniew et al., 2016); 4) calcium imaging can also be combined with optogenetic manipulations, which makes it possible to perform all-optical probing of circuit function (Packer et al., 2015); 5) recordings can be performed in freely moving animals, providing a key link between circuit activity and behavior (Lin and Schnitzer, 2016); and 6) calcium imaging is less invasive than traditional electrode recordings (e.g., tetrodes, silicon probes). Another advantage the technique offers is the ability to precisely map the relative spatial location of groups of neurons. At the same time, lower costs of lasers and microscope components, including commercial systems, are making it increasingly affordable for laboratories to adopt calcium imaging.

Analyzing data generated by fluorescence calcium imaging, however, is not trivial. While increasing numbers of scientists can now afford to purchase a 2-photon microscope to record with the latest genetically encoded fluorescent calcium indicators (Chen et al., 2013), they often lack the expertise to write the required sophisticated software to analyze the dynamic signals. The scientists who adopted 2-photon microscopy two decades ago had to write their own custom code. Unfortunately, it is often the case that the custom routines developed by one laboratory cannot easily be incorporated by other groups because the data are acquired differently or because the code may appear cryptic to those inexperienced in programming. Although there have been some recent efforts to develop and distribute open-source software packages for extracting and analyzing dynamic fluorescent signals, such as *CaImAn*, *SIMA*, and *Suite2P* (Giovannucci et al., 2019; Kaifosh et al., 2014; Pachitariu et al., 2017), these still require some programming experience in MATLAB or Python. Consequently, the lack of a simple, easy to use resource for analysis of calcium signals limits access to these powerful techniques for mainstream neuroscience laboratories and also affects the replicability of experiments, both of which are vital to scientific progress.

To address this unmet need, we created a new software package for analysis of calcium imaging data that emulates similar tools for analysis of volumetric or functional MRI data or for spike sorting in electrophysiology (Chung et al., 2017; Yger et al., 2018). Our goal was to automate all of the core aspects of calcium imaging analysis in a modular design: 1. image registration (motion correction); 2. image segmentation into distinct regions of interest (ROI detection and refinement); 3. signal extraction and deconvolution (dimension reduction). The entire process can be fully automated (e.g., unbiased ROI selection), allows for batch processing, and is compatible with standard desktop or laptop computers. The code is well-documented with discrete variable and function names to allow users to examine how the code works and make modifications. For example, it allows users to implement new routines or remove existing functionalities. Importantly, the code is wrapped in a set of user-friendly and intuitive graphical user interfaces (GUIs), which permits scientists who are not yet proficient in writing code in MATLAB to quickly and easily work with the interface and modify it as needed. This toolbox, *EZcalcium*, accepts common standards of data formats and should therefore be compatible with a variety of microscope systems. For motion correction we incorporated *NoRMCorre* because it achieves excellent results with a variety of recordings (Pnevmatikakis and Giovannucci, 2017). For the segmentation process, we incorporated aspects of *CaImAn* (Giovannucci et al., 2019), which is able to detect ROIs dynamically, but we included manual selection capabilities as well. Finally, *EZcalcium* includes an ROI refinement step that allows the user to inspect the traces and the shape of each ROI and set parameters to automatically or manually remove ROIs.

The toolbox is efficient and fast, such that even standard desktop computers can process multi-gigabyte video files in a few hours. As far as signal extraction and analysis, the code uses established constrained non-negative matrix factorization algorithms (Pnevmatikakis et al., 2016) to predict which changes in fluorescence intensity correspond to biological signals (action potentials) and which represent noise. Most importantly, the software is open-source and freely available on GitHub, so that future users can then contribute to its enhancement and expansion.

## 2 Method

### 2.1 Equipment

Programming experience is not required to run the toolbox. We recommend MATLAB R2018b (or newer MATLAB) on any operating system for using the *EZcalcium* toolbox within MATLAB. The toolbox was finalized and tested heavily in R2018b (9.5) and is likely to be the most compatible without modification in that environment. A 64-bit version of MATLAB running on a 64-bit computer is required to process files over 800 MB. The following MATLAB toolboxes are also required when running the source scripts: Signal Processing, Statistics, and Parallel Computing. The amount of available system RAM necessary for a system depends on the size of the data being processed. Ideally, the amount of system RAM should be at least 3X the file size of raw, uncompressed data. *Motion Correction* is the most RAM-intensive step of the process and significant slowdowns may occur if the necessary RAM is not available. CPU requirements for the toolbox are minimal, but processing is vastly improved with multiple cores. The toolbox also runs faster when the data to be analyzed is located on a fast, solid-state hard drive, since large amounts of data must be read and, in the case of *Motion Correction*, also written.

### 2.2 Software setup

*EZcalcium* is freely available to be downloaded from the Portera-Cailliau Lab website (https://porteralab.dgsom.ucla.edu/pages/matlab) or its GitHub site (https://github.com/porteralab/EZcalcium). For running *EZcalcium*, MATLAB 2018a or newer is required. To run the toolbox source code, first add the toolbox folder to the MATLAB path directory. Type the command *EZcalcium*. A GUI will load that can run all the individual modules. When working in Windows, data saved to directories listed under C:\Users, C:\Program Files, and the directory in which MATLAB is installed are often protected against writing and deletion. Therefore, in order to process and generate data, it is recommended to use imaging files saved outside of C:\Users, C:\Program Files, and the directory in which MATLAB is installed. Failure to do so may result in an error stating “You do not have write permission.”

### 2.3 Calcium imaging

We provide examples of typical results from using *EZcalcium* to analyze two-photon calcium imaging data acquired in drosophila and mice. The mouse data was collected in the Portera-Cailliau lab. Viral vectors expressing GCaMP6s (rAAV-hSyn-GCaMP6s) were injected in the VPM nucleus of the thalamus or in primary visual cortex at the time of the cranial window surgery, as previously described (Goel et al., 2018). Cranial windows were implanted over the barrel cortex (thalamocortical axons in layer 4) or primary visual cortex (layer 2/3 neurons), as described (Holtmaat et al., 2009). After allowing 2-3 weeks for optical expression of the indicator, imaging sessions began using custom-built 2-photon microscopes with a resonant mirror at 30 Hz (visual cortex) or galvo mirrors at 8 Hz (axon boutons). These experiments followed US National Institutes of Health guidelines for animal research, under an animal use protocol (ARC #2007-035) approved by the Chancellor’s Animal Research Committee and Office for Animal Research Oversight at the University of California, Los Angeles.

Drosophila two-photon calcium imaging data from was obtained from R8 photoreceptors in the proximal medulla of the senseless-FLP fly line (UAS-opGCaMP6s/GMR-FRT-STOP-FRT-Gal4; UAS-opGCaMP6s/UAS-10X-myr::tdTomato), as described (Akin et al., 2019). Images were collected at 1 frame every 15 s.

## 3 Results

### 3.1 Overview

The *EZcalcium* toolbox is controlled by a set of intuitive and user-friendly GUIs. Once configured for a set of similarly acquired data, preferences can be saved and reused for simple automation of workflow. The toolbox is able to import a variety of imaging data file types (including .tif, .avi, .mat) and supports the export of data in both proprietary (.mat, .xlsx) and open (.csv) file formats.

*EZcalcium* contains three main modules: *Motion Correction*, *ROI Detection*, and *ROI Refinement*. (Fig. 1). The *Motion Correction* module consists of a non-rigid method of template matching, *NoRMCorre* (Pnevmatikakis and Giovannucci, 2017). The *ROI Detection* module, which is built off of the *CaImAn* toolbox (Giovannucci et al., 2019), includes automated ROI detection, signal extraction, and deconvolution of fluorescence calcium signals. The *ROI Refinement* module enables the user to sort and view ROIs, manually exclude ROIs, and use automated and customized ROI exclusion criteria, including spatial and activity-dependent metrics, to refine the list of ROIs that will be ultimately analyzed.

**Figure 1.**
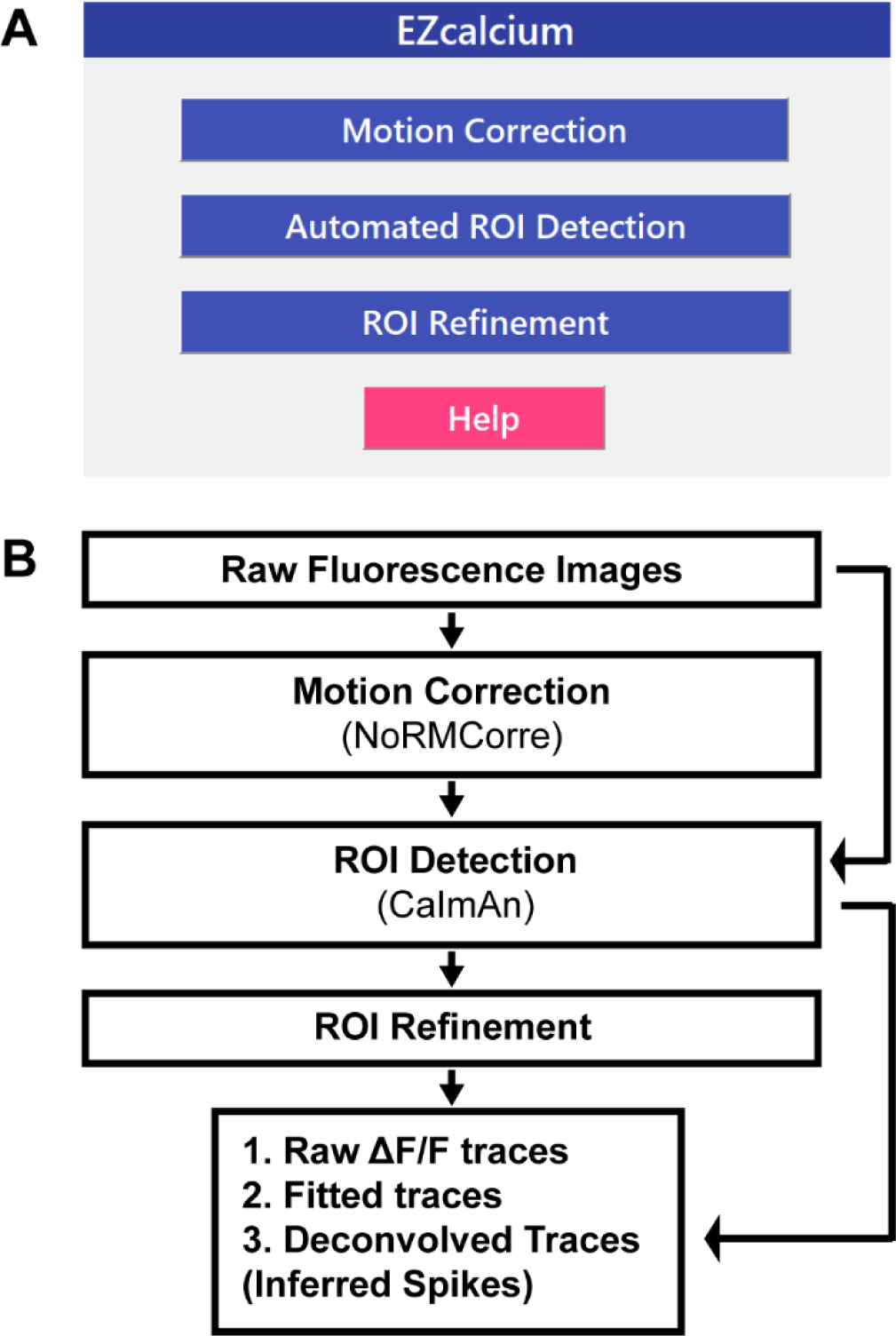
EZcalcium control GUI and workflow. A. Appearance of the main *EZcalcium* control GUI. Each of the 3 modules of the toolbox is called by clicking on its respective button. Help buttons are located throughout the toolbox to provide documentation and resources to the user. B. Workflow of the *EZcalcium* toolbox.

An important feature of *EZcalcium* is that batch processing is supported in the most resource-intensive modules (*Motion Correction* and especially *ROI Detection*), which allows for a large number of files to be sequentially loaded, processed, and saved. This feature is ideal for processing large data sets (> 4 GB) that would best be processed overnight or on a dedicated analysis machine.

In the event of a system ‘crash’ or shutdown, as can occur during an automatic operating system update, progress is automatically saved at the end of each successful file processing and can be resumed from the start of the previous incomplete attempt.

The MATLAB source code is freely available for all modules so that users may modify and adapt the code for their own purposes (https://github.com/porteralab/EZCalcium). It is well-documented within the code itself for easy understanding and to facilitate modification. The toolbox is designed to be modular and adaptable for use with dynamic fluorescence data obtained from a wide variety of imaging systems; it should easily translate to other types of dynamic fluorescence imaging, such as voltage sensors and indicators of neurotransmitter release.

We have tested the *EZcalcium* toolbox on calcium imaging data collected with in vivo two-photon microscopy in neurons expressing GCaMP6s in mouse barrel cortex and primary visual cortex, in the visual system of Drosophila larvae, in boutons of thalamocortical axons in layer 4 in mouse barrel cortex, or in cortical astrocytes. Examples of such analyses are provided throughout.

Below, we describe each of the three independent modules of *EZcalcium* (*Motion Correction*, *ROI Detection*, and *ROI Refinement*) and illustrate the end result with specific examples from in vivo 2-photon calcium imaging experiments done in mice and Drosophila. A more detailed step-by-step walkthrough tutorial is presented in the Supplementary Materials, EZcalcium Procedures.

### 3.2 Motion Correction

Motion correction is often needed when imaging in awake behaving animals, but it can be challenging to use some of the more popular approaches (e.g., TurboReg in ImageJ) for image registration, due to the nature of dynamic fluorescence signals. The main problem is that, as animals move, there is primarily translational drift in x and y directions, which is relatively easy to correct. However, the fluorescence signal intensity of the profile being imaged (cell body, axon bouton) will fluctuate depending on whether the element is active or not, which simple static registration algorithms may mistake for actual translation in the x-y dimension. The most popular alignment algorithms have been those based on template matching strategies, such as the Lucas-Kanade (Greenberg and Kerr, 2009; Mineault et al., 2016) or the Non-Rigid Motion Correction (*NoRMcorre*) strategy (Pnevmatikakis and Giovannucci, 2017). For *EZcalcium* we implemented the *NoRMcorre* motion correction algorithm (Fig. 2) because it was significantly faster than Lucas-Kanade and a single iteration was sufficient to achieve satisfactory registration (Pnevmatikakis and Giovannucci, 2017). We found that this motion correction procedure provides excellent results for 2-photon microscopy data collected from awake behaving mice (cell bodies or axon boutons) or live Drosophila visual system (Fig. 3). The motion corrected video is saved in TIFF format. Of course, it is not strictly necessary to use the *Motion Correction* module if the videos do not exhibit motion artifacts (e.g., calcium imaging in brain slices or anesthetized animals).

**Figure 2.**
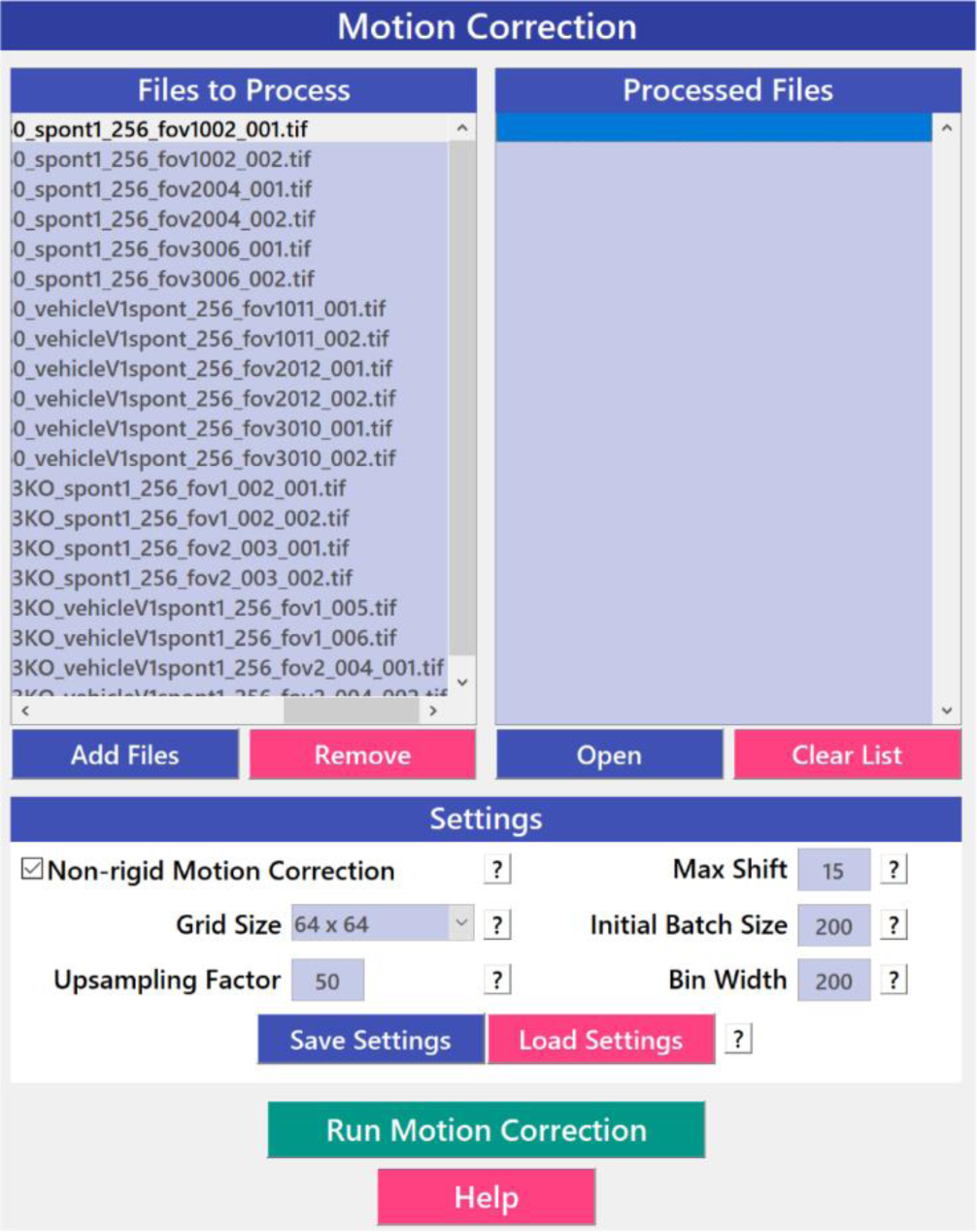
Motion Correction GUI. The *Motion Correction* module utilizes the *NoRMcorre* algorithm to align the image sequences. It allows for batch processing of multiple videos.

**Figure 3.**
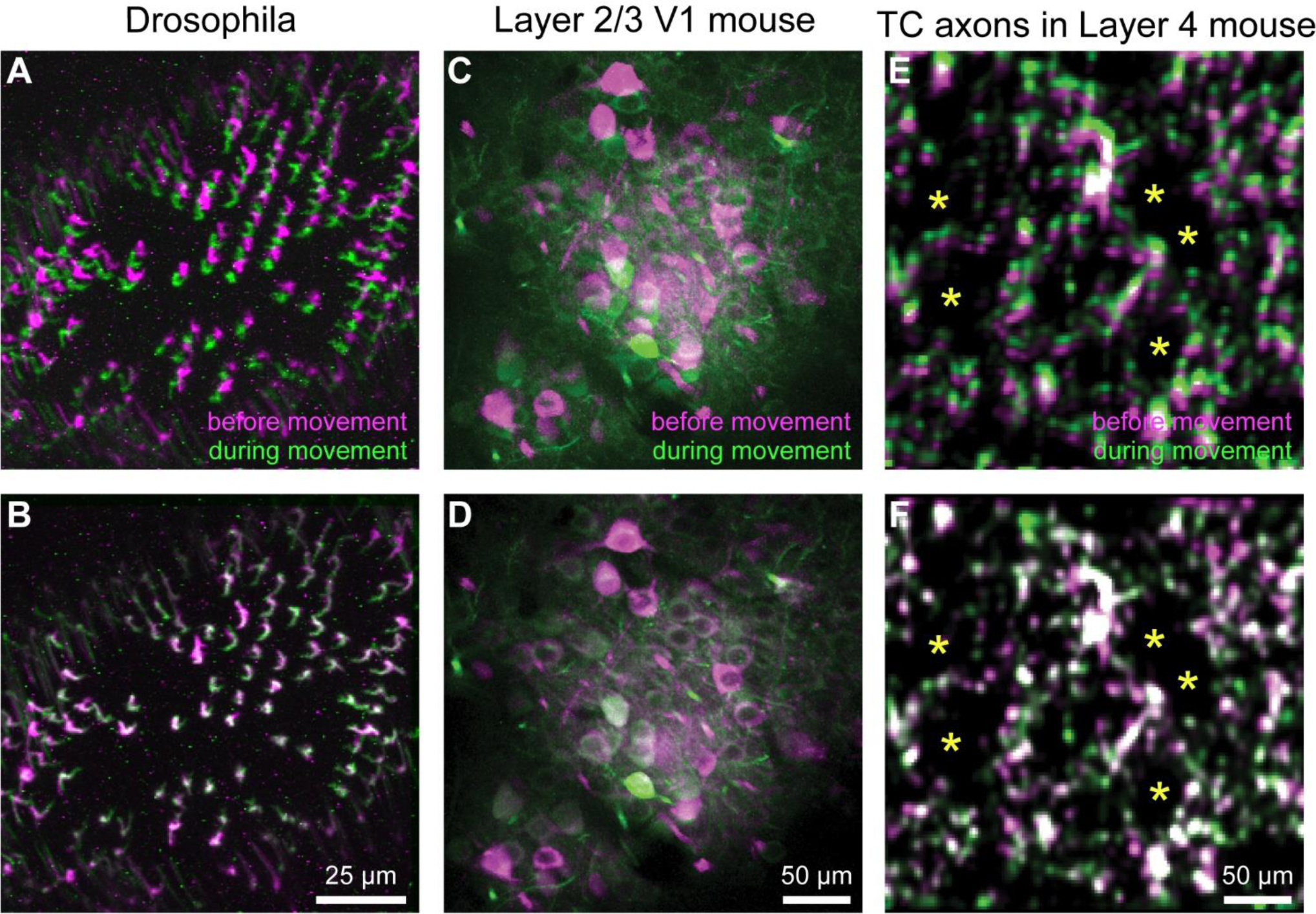
Motion Correction Results. A-F. Representative xyt MAX intensity projections before (A, C, E) and after image registration (B, D, F) with *EZcalcium* for calcium imaging recordings of R8 photoreceptors in the proximal medulla of drosophila larvae (A, B), layer 2/3 neurons in adult mouse visual cortex (C, D) and thalamocortical axon boutons in layer 4 of adult mouse barrel cortex (E, F). Purple and green represent maximum intensity projections of 100-150 frames before and during motion artifact.

### 3.3 ROI Detection

After image sequences (‘videos’) are motion-corrected, the next step in processing the data is to segment the image into desired ROIs with the *ROI Detection* module (Fig. 4). With the large fields of view typically collected with calcium imaging, manual segmentation of the images into distinct ROIs can be time consuming and impractical, and potentially introduces human error, such as biased exclusion of ROIs that are smaller or less active. It is also challenging to separate signals from juxtaposed or overlapping ROIs. Automation of ROI segmentation can diminish the effects of bias and significantly reduce the time required to process large data sets. Our *ROI Detection* module is based on the *CaImAn* toolbox (Giovannucci et al., 2019), and provides access to most of the basic functionalities provided by *CaImAn*. Segmentation in *EZcalcium* with the *ROI Detection* GUI allows for a manual initial refinement step as an option for adding or removing ROIs by hand (Fig. 5A, D). When such manual refinement is not selected, batch processing proceeds in a fully automated fashion. *ROI Detection* is based on using both temporal and spatial correlations to identify nearby pixels that exhibit similar changes in fluorescence intensity at the same time. It initializes by estimating activity using power spectral density or sparse non-negative matrix factorization (Maruyama et al., 2014). The *ROI Detection* step generates a MATLAB data file containing extracted fluorescence (ΔF/F) and deconvolved neural activity (spiking) traces using sparse non-negative deconvolution algorithms for each ROI (Giovannucci et al., 2019; Pnevmatikakis et al., 2016; Vogelstein et al., 2010). Representative examples of segmented fields of view of neuron somata in mouse visual cortex, R8 photoceptors in Drosophila and axon boutons in mouse barrel cortex are provided (Fig. 5).

**Figure 4.**
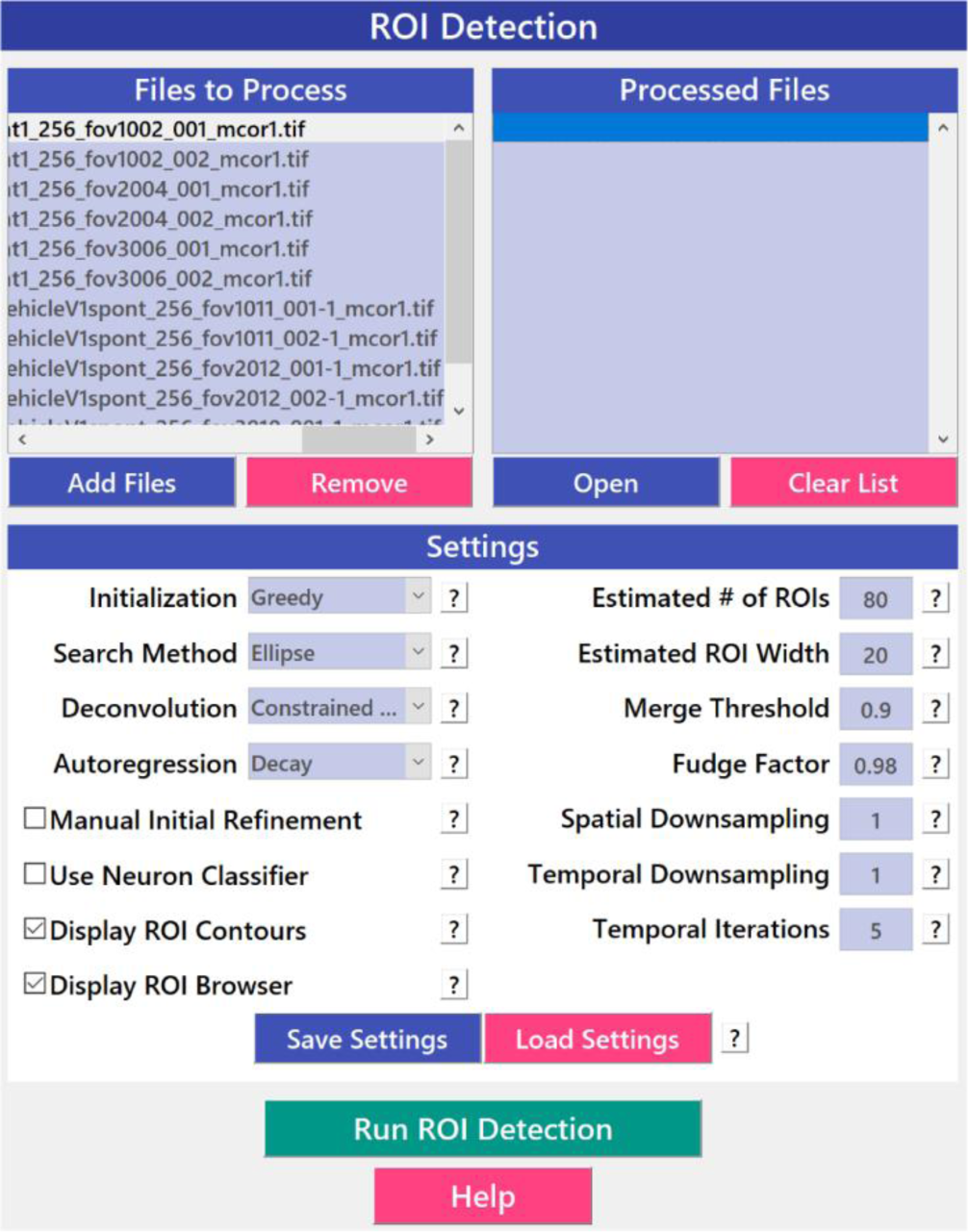
ROI Detection GUI. The automated *ROI Detection* module is used to segment the image files into individual ROIs. It minimizes the impact human bias in ROI selection and allows for batch processing of multiple videos. ROIs that are not automatically detected can be manually added.

**Figure 5.**
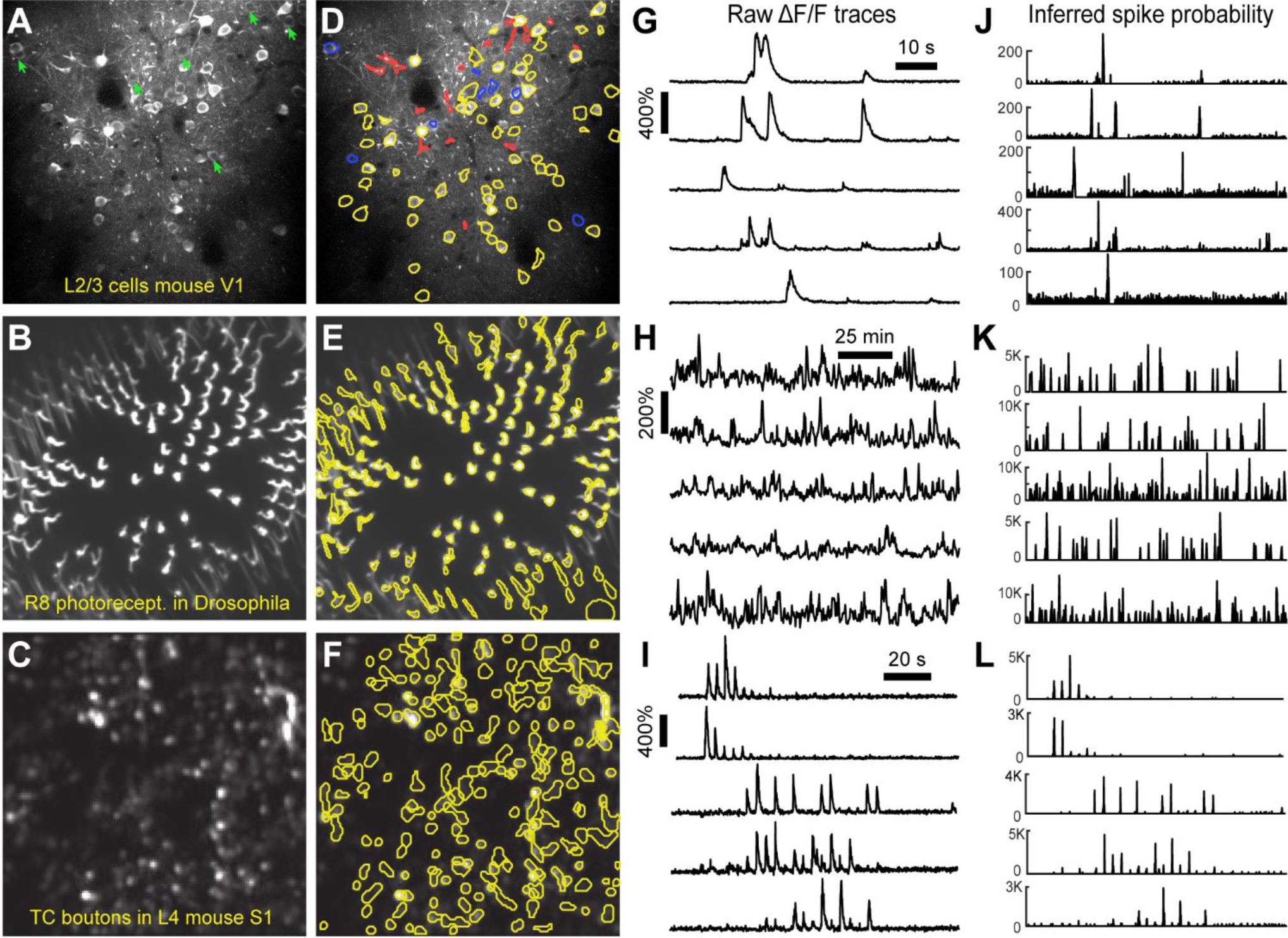
ROI Detection Results. A, B, C. Representative examples of a field of view from two-photon calcium imaging of layer 2/3 neurons in visual cortex of adult awake mice (A), R8 photoreceptors in the proximal medulla of drosophila larvae (B), and thalamocortical axon boutons in layer 4 of barrel cortex in adult anesthetized mice (C). Green arrows point to inactive cells that were not automatically detected and could not be manually added. D, E, F. Same fields of view after automated ROI detection. Red contours indicate ROIs that might be removed during the *ROI Refinement* module due to their shape. Blue contours indicate ROIs that were selected manually. Yellow and red contours are ROIs that were selected automatically. G, H, I. ΔF/F traces of representative ROIs from the same experiments as in D, E and F. The traces in G and H show spontaneous activity, whereas traces in I represent whisker-evoked activity (20 stimulations for 1 s at 10 Hz, with 3 s long inter stimulus interval). J, K, L. Inferred probability of spiking after deconvolution for the same ROIs as in D, E and F.

Multiple temporal iterations can be used to improve accuracy and to detect ROIs of complex shapes, which could be relevant for irregular profiles such as those of glia, invertebrate neurons, or dendritic spines. After processing a single video file, the user can find measurements of actual ROI size in the next module, *ROI Refinement* (see below), which can then be used to improve the performance of further ROI detection.

### 3.4 ROI Refinement

Automated ROI detection algorithms are still not perfect, and users may find it necessary to manually inspect shapes and individual traces of detected ROIs in order to reject false positive ROIs. The *ROI Refinement* GUI in *EZcalcium* (Fig. 6) automates the exclusion of certain ROIs using user-defined heuristics. By automating this process, operator biases can be minimized. Refinement criteria include characteristics of ROI morphology and activity. The *ROI Refinement* module also allows the user to readily view the characteristics of all detected ROIs, including a visual map of the location of each ROI, the shape of the isolated ROI, and the extracted fluorescence traces in raw, inferred, and deconvolved formats (Fig. 7A). This helps define what criteria should be used for excluding ROIs from further analysis. It can also be used to manually exclude ROIs. After the *ROI Refinement* step is completed, data can be exported into proprietary (.mat, .xlsx) or open file formats (.csv).

**Figure 6.**
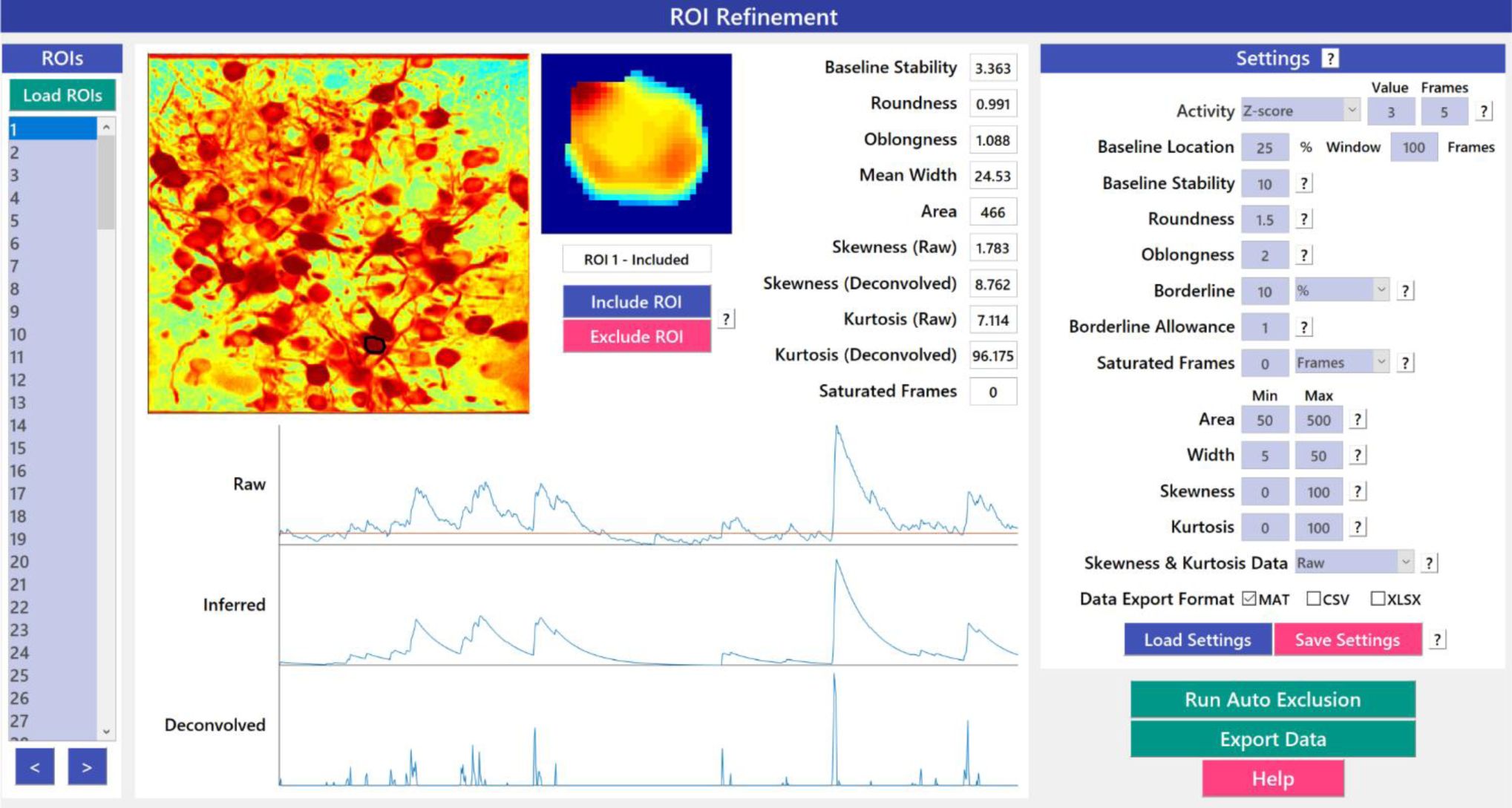
ROI Refinement GUI. *ROI Refinement* is the final step of the *EZcalcium* workflow and consists of automatically or manually excluding ROIs based on their shape or their traces. Following refinement, data can be exported in a variety of formats.

**Figure 7.**
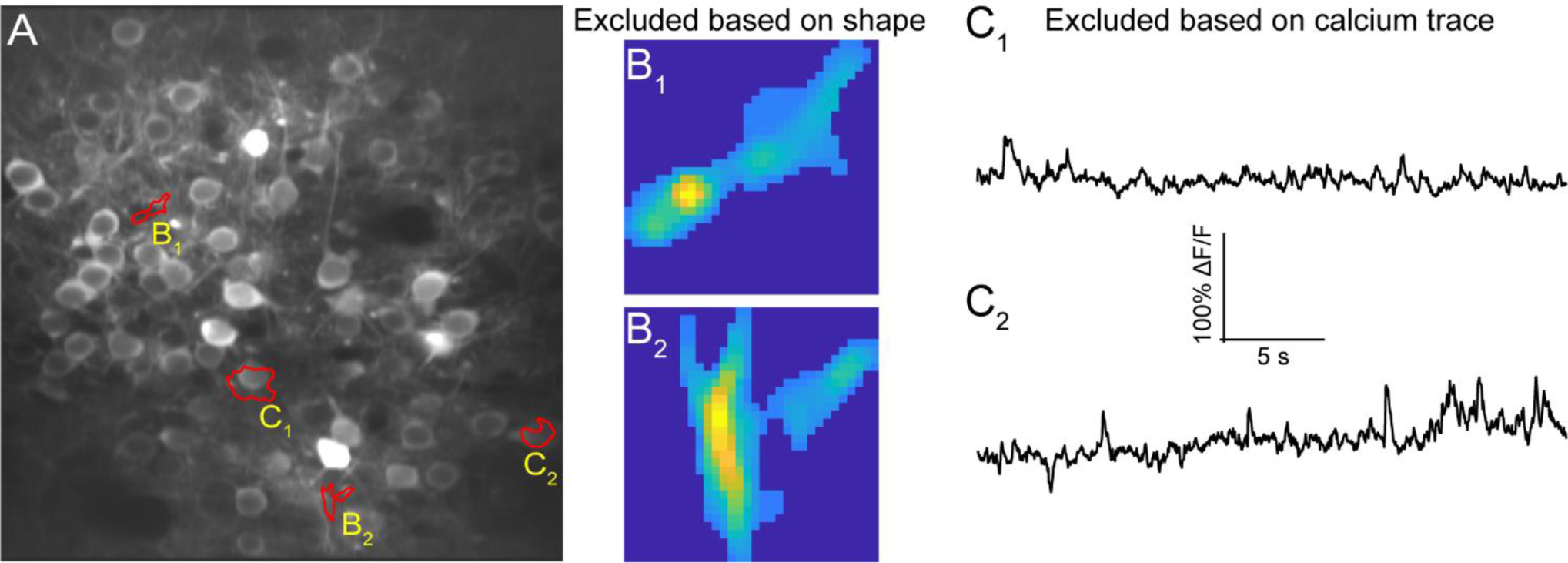
ROI Refinement Results. A. Representative field of view from a calcium imaging experiment of layer 2/3 neurons in visual cortex in an adult mouse (GCaMP6s) showing ROIs that showed unusual shapes or traces. B1, B2. Representative ROIs to be excluded based on their shape. C1, C2. Representative ROIs to be excluded based on their calcium traces.

## 4 Discussion

Despite its rising popularity in neuroscience, analyzing the rich data generated by fluorescence calcium imaging can be challenging. The goal of this project was to develop an open source toolbox for analysis of calcium imaging data that could be freely disseminated across the broader scientific community. Aspects of certain modules of the *EZcalcium* toolbox have already been successfully implemented in several published studies (Goel et al., 2018; He et al., 2018; He et al., 2017) and the toolbox is freely available on GitHub (https://github.com/porteralab/EZcalcium).

Our main priority was to create a toolbox with simple and intuitive GUIs that provide a ‘push-button’ feel for the user. *EZcalcium* is well documented and includes discreet variable and function names that allow users to examine how the code works and to make modifications, as well as help features for every button. We also wanted *EZcalcium* to be comprehensive, meaning that it would accomplish the main steps that most users would require: namely, image registration (*Motion Correction* module), segmentation (*ROI Detection*), and signal extraction (*ROI Refinement*). We therefore incorporated the latest MATLAB resources for analysis of calcium imaging data (Giovannucci et al., 2019; Pnevmatikakis and Giovannucci, 2017; Pnevmatikakis et al., 2016; Vogelstein et al., 2010).

*EZcalcium* is designed for those investigators who may find *CaImAn* less intuitive, or for those who prefer to use MATLAB rather than Python for their customized changes to the code. In addition, the *ROI Refinement* module in *EZcalcium* can be used as a browser to inspect all detected ROIs, and to eliminate false-positive ROIs. We consider this to be a critical step because the automatic *ROI Detection* step is still prone to false positives and there are advantages to having additional manual control in omitting certain ROIs, after inspecting their raw traces within the GUI.

All three fundamental steps in the processing of dynamic fluorescence signals in biology (*Motion Correction*, *ROI Detection*, and *ROI Refinement*) are major components of *EZcalcium*. Although we designed *EZcalcium* primarily with two-photon calcium imaging in mind, the toolbox should in principle be compatible with other types of dynamic fluorescence data, such as voltage indicators (Abdelfattah et al., 2019) and fluorescence sensors that probe neurotransmitter release (Marvin et al., 2013). The open source and modular format allow our toolbox to be readily adapted for specific needs so different users can add new capabilities. For example, while the *ROI Detection* module in *EZcalcium* is presently not compatible with microendoscopic calcium imaging data (miniscope), an additional module with Constrained Non-negative Matrix Factorization for endoscopes [CNMF-E] (Zhou et al., 2018) could easily be added to our toolbox to segment those data.

## Supporting information

EZcalcium procedures

## 6 Author Contributions

DC and CP designed the framework of the toolbox. DC, BW, MG wrote the code of the toolbox. BW, CH, AG, AS, NK, EA, WZ provided experimental data and tested the toolbox. DC, BW, MG, CP wrote the manuscript.

## 8 Acknowledgments

We thank Lauren Wierenga for her help in earlier stages of this project, Bryce Bajar, Orkun Akin and Larry Zipursky for the Drosophila data, as well as Dean Buonomano and Dario Ringach for feedback on the manuscript. Some aspects of the *EZcalcium* code were described in Dr. Daniel Cantu’s PhD thesis: *Analysis of fluorescent calcium signals in the detection of neural circuity abnormalities in a mouse model of autism* (https://escholarship.org/uc/item/48s773xv) (Cantu, 2019).

