## Supplementary material for "EZcalcium: Open Source Toolbox for Analysis of Calcium Imaging Data": EZcalcium procedures

Below are step-by-step instructions for using the three modules of *EZcalcium* toolbox.

### Motion Correction

1. From the initial GUI, click on the box titled *Motion Correction* to load the *Motion Correction* GUI (Fig. 2).
2. In the *Motion Correction* GUI, click the Load button to choose any number of videos you would like to load. You can drag a box or use the select or shift key to select multiple files with the mouse. If you close the file select window and decide to add more files later (e.g., from a separate folder), they will appear at the bottom of the list. If you accidentally try to add the same file multiple times, it will not be added to the list again and an error will display. Compatible file types include multipage 8- or 16-bit .tif, .avi. Files used need to be single-channel images. .tif files greater than 4 GB need to be in an uncompressed format. If your data is of an incompatible file type, it can likely be converted to .tif or .avi with free software such as ImageJ, available at https://fiji.sc/.
3. To remove a file that has been added to the Files to Process list, select the file with the mouse and click the Remove button.
4. Choose whether to use non-rigid motion correction or not. Non-rigid Motion Correction splits the field of view into a number of overlapping patches to correct for within-frame motion artifacts. While this can give more accurate results, it is considerably slower than rigid motion correction.
5. Grid Size defines the size of each patch in pixels when using non-rigid motion correction. It will have no effect if Non-rigid Motion Correction is not selected.
6. Upsampling Factor defines the upsampling factor for subpixel registration.
7. Max Shift defines the maximum rigid shift in pixels allowed in each direction.
8. Initial Batch Size defines the number of frames from the beginning used for calculating the initial template.
9. Bin Width defines the number of frames of each bin, over which the registered frames are averaged to update the template.
10. The Save Settings button allows the user to save all settings under a specific name of one's choosing. Settings are saved as .mat files. These include all settings in the Settings section.
11. The Load Settings button allows one to load all saved settings in future sessions.
12. Once all settings are configured, click the Run Motion Correction button. Progress of motion correction will be displayed in MATLAB command window.
13. After motion correction is completed, the new file generated will be added to the bottom of the Processed Files list. Original files will not be overwritten. A new file will be created with _mcor appended to the filename.
14. The Open button will open the selected file in the default program set by the operating system.
15. Clear List clears the entire list of Processed Files.

### ROI Detection

1. From the initial GUI, click on the icon titled Automated ROI Detection to load the *ROI Detection* GUI (Fig. 4).

2. In the *ROI Detection* GUI, click the Load button to choose any number of videos you would like to load. Compatible file types include multipage .tif, .avi. The user closes the file select window and decides to add more files later, such as from a separate folder, the files will appear at the bottom of the list. If the user accidentally tries to add the same file multiple times, it will not be added to the list again and an error will display. File type requirements for *ROI Detection* are similar to those of *Motion Correction*. When troubleshooting and testing motion correction for the first time, it is advisable to start with correcting a single file and testing the settings.
3. To remove a file that has been added to the Files to Process list, select the file with the mouse and click the Remove button.
4. Initialization methods for providing an initial estimate of spatial and temporal components: Greedy is recommended for videos of neuronal somata. It relies heavily on spatial components and generally runs much faster; Sparse NMF is recommended for more complex structures, such as dendrites, dendritic spines, or axons.
5. Search Method determines the spatial components (location) of ROIs: Ellipse assumes components have an ellipsoid shape, such as for neuronal somata; Dilate can be used with either ellipsoid or non-ellipsoid ROIs, but generally takes longer.
6. Deconvolution determines the method for translating activity-induced changes in the fluorescence intensity of the indicator into approximate firing rates. If you are imaging an organism that does not produce action potentials, set this to the fastest setting available. Constrained FOOPSI is generally recommended and, with CVX, it is typically the fastest method of deconvolution when working with high signal-to-noise traces. Constrained FOOPSI – SPGL1 works better with medium-to-low signal-to-noise traces. MCMC is a fully-Bayesian deconvolution method that is computationally intensive and is recommended when higher precision is required. MCEM alternates between the listed Constrained FOOPSI deconvolution and MCMC to update time constants, which is significantly faster than MCMC alone. Note that additional installation of CVX is needed when using MCMC. Users can download CVX at [http://cvxr.com/cvx/download/](http://cvxr.com/cvx/download/), and follow the instructions.
7. Autoregression is used to estimate the calcium indicator kinetics. Rise and Decay estimates both the rise and decay kinetics of the calcium indicator and incorporates them when extracting fluorescence traces and deconvolving the signal. Due to the difficulty in detecting fast rise times, using Rise and Decay may result in overfit data if the imaging was performed at low temporal resolution (< 16 Hz). Decay estimates only the decay kinetics of the calcium indicator and is the recommended setting for lower temporal resolution imaging. No Dynamics will produce only raw traces and will not perform deconvolution.
8. Manual Initial Refinement adds an additional step following initialization to manually add or remove ROIs. ROIs can also be removed in the step *ROI Refinement*. To fully automate the process, it is recommended to optimize your settings to slightly overestimate the number of ROIs and then remove erroneous ROIs later. This step is included as an option for particularly troublesome files and for those who prefer semi-automated ROI selection. Initial spatial components will be displayed in a new figure. The center of estimates in ROIs is highlighted with a magenta circle, surrounded by the boundary of the ROI. To manually add an ROI, left click with the mouse where you want to add the center of an ROI. The boundary of the ROI will be automatically computed and drawn. To manually remove an ROI, right click on any ROI center. Hit the ENTER/RETURN key to continue ROI detection.
9. Use Neuron Classifier utilizes *CaImAn*'s built-in neuron classifier to exclude potentially erroneous ROIs based on their shapes. Leave this option unselected if desired ROIs are not neuron cell bodies.

10. Choose which additional figures you want generated. When batch processing many files, it is not recommended to use these. Display ROI Contours generates a map with all the ROI boundaries, each labeled with the same ROI number as was used in the data. Display ROI Browser shows extracted raw fluorescence data, the inferred trace generated, and the ROI shape and location.
11. Estimated # of ROIs is the estimated maximum number of components in the field of view. This must be set to a minimum value of 1. If you want to perform fully manual ROI selection, set Estimated Components to 1 and check the box to enable Manual Initial Refinement. When the manual refinement step starts, delete the initial automatically detected ROI. If no components are determined to be similar enough to be merged, the most likely result is that the number set for Estimated Components is the number that will be initially detected.
12. Estimated ROI Width is the estimated width, in pixels, of your components. If you have a simple ROI shape, such as a cell soma, you can use the width of the entire ROI as your component width.
13. Merge Threshold is the threshold at which two components will be merged into a single ROI. Components that share a correlation coefficient above Merge Threshold will be merged into a single ROI.
14. Fudge Factor is useful for estimating time constants of very noisy data, in particular those with low temporal resolution (slow frame rate). The value indicates a multiplicative bias correction for time constants of each ROI during deconvolution. Fudge Factor should generally be set to 0.95-1. A value of 1 indicates that no bias correction will be performed.
15. Spatial Downsampling will downsample the spatial resolution of a video by a factor set here. The value entered should be a positive integer. A value of 1 means that no downsampling will be performed. This is useful for rapidly troubleshooting the settings on videos with a very large field of view.
16. Temporal Downsampling is similar to Spatial Downsampling, except it downsamples the temporal resolution. This is useful for optimizing settings on very long, high frame rate (>15 Hz) videos.
17. Temporal Iterations is the number of iterations that will be performed to calculate the temporal components of ROIs. This should be set to at least 2 when not rapidly testing other parameters.
18. Save Settings allows you to save all settings under a specific name of your choosing. Settings are saved as .mat files.
19. Load Settings allows you to load any previously saved settings.
20. Once all settings are configured, click the Run ROI Detection button.
21. After ROI detection is complete, the new file generated will be added to the bottom of the Processed Files list. A .mat file will be created with the complete set of data generated during The Open button will open the selected .mat file into a MATLAB workspace, if running source scripts.
22. Clear List clears the entire list of Processed Files.

**ROI Refinement**

1. From the initial GUI, click on the icon titled *ROI Refinement* to load the *ROI Refinement* GUI (Fig. 6). In the *ROI Refinement* GUI, click the Load ROIs button to choose a data file generated by *ROI Detection* you would like to load.

2. Select an ROI by clicking on it to view. The left arrow button (<) will automatically select and load the previous ROI (lower in ROI number). The right arrow button (>) will automatically select and load the next ROI (higher in ROI number).
3. The information of the currently selected ROI is shown in the middle panel of the GUI. The whole field of view is displayed next to the ROI list, with the current ROI contour highlighted. The enlarged view of the current ROI is shown on the right. Below the ROI figures are the extracted traces, including fluorescence and deconvolved traces.
4. Settings: Activity can be set to include ROIs that surpass a chosen activity threshold for a given number of consecutive frames. This threshold can be set in the Value box in units of Z-Score, and the chosen number of consecutive frames can be set in the Frames box. Baseline Location determines the percentage of the data from the beginning and end of the recording that will be considered when determining the baseline activity level; for example, if this is set to 25%, the first 25% and the last 25% of the total frames will be used to determine the baseline values. *EZcalcium* will find the minimum value within these frames and use the surrounding frames to calculate a mean baseline value. Window determines the number of frames to be averaged to find this baseline value. Baseline Stability is used to check if an ROI has a stable baseline throughout the imaging session by comparing the baseline at the beginning of recording with the baseline at the end. Roundness measures how similar an ROI is to a circle. This is useful when you are looking exclusively for neuron somata or other round ROIs. Oblongness is a measure of the ellipsoid shape of an ROI. It is calculated by dividing the length of the major axis of the ellipse by the length of the minor axis. This is useful in excluding overly oblong ROIs that are not likely to be neurons.
5. Borderline % can be set to allow an ROI to have criteria slightly outside of the desired range. If an ROI has a number of criteria within the Borderline range equal to or less than the Borderline Allowance, the ROI will be included.
6. Saturated Frames can be used to set a maximum number of frames in which the ROI can be fully saturated. This can also be input in terms of the percentage of total frames.
7. The next section allows the user to set minimum and maximum values for a variety of parameters. Area measures the area of an ROI and is useful for excluding ROIs that are too small or too large. Width indicates the mean width of an ROI and the relative size of the ROI. It is useful for removing ROIs that result from having too many components coalesce together into a single profile. Skewness and Kurtosis are used to describe the probability distribution of either the dF/F data or the deconvolved data for an ROI. Skewness measures the left- or right-skewness of the data. Kurtosis measures the likelihood of finding outliers in a distribution. For example, a cell with large, fast, and frequent spikes in calcium levels will have a higher kurtosis value than cells with a more moderate distribution of slow activity. Skewness & Kurtosis Data is used to select whether the dF/F or deconvolved data will be used as the basis of the exclusion criteria for these two parameters.
8. Click the Run Auto exclusion button to automatically exclude ROIs based on your chosen criteria.
9. To manually exclude an ROI, select an ROI and click the Exclude ROI button. After being excluded, the ROI number in the list will be stricken out and the Exclude ROI button will become an Include ROI button.
10. Click the Export Data button to export your data in your selected format.
